# Modular gene interactions drive modular pan-genome evolution in bacteria

**DOI:** 10.1101/2022.11.15.516554

**Authors:** Juan C. Castro, Sam P. Brown

## Abstract

Depending on the scale of observation, bacterial genomes are both organized and fluid. While individual bacterial genomes show signatures of organization (e.g., operons), pan-genomes reveal genome fluidity, both in terms of gene content and order (synteny). Here we ask how mutational forces (including recombination and horizontal gene transfer) combine with selection and gene interactions to shape genome organization and variation both within and across strains. We first build an evolutionary simulation model to assess the impact of gene interactions on pan-genome structure. A neutral evolutionary model can produce transient co-segregation of initially linked genes but is vulnerable on longer time-scales to perturbing mutational events. In contrast, incorporation of modular gene fitness interactions can produce sustainable clusters of linked and co-segregating genes, with the network of co-segregation recapitulating the defined simulation ‘ground-truth’ network of gene interactions. To test our model predictions, we exploit the increasing number of closed genomes in model species to define gene co-segregation networks in the pan-genomes of *Escherichia coli* and *Pseudomonas aeruginosa*. Using these highly curated pan-genomes, we identify modular clusters of physically linked and co-segregating genes and show that the resulting co-segregation networks map onto underlying gene-regulatory and metabolic gene interaction networks. The results imply that co-segregation networks can contribute to accessory genome annotation, and more generally that gene interactions are the primary force shaping genome structure and operon evolution.

## Introduction

Individual bacterial genomes show hallmarks of organization (1). For example, genes involved in related biochemical processes are often clustered together on the chromosome (2). The organization of clustered genes can be refined to the point where they are transcribed together as operons, with a shared promoter and operator sequences. Regulatory genes are also often found close to the genes they regulate. A classic example is the *lacI* repressor gene, which resides near the *lacZYA* operon in *Escherichia coli* (3).

While organized, genome structure is under constant assault from an array of mutational processes that scramble gene content and order. Genes can be gained or lost. Whole sequences of genes can be flipped (inversions), moved along a chromosome (translocation, recombination) or shuttled to a new individual (horizontal gene transfer). As a result of these perturbing forces, genomic organization is variable across bacterial isolates; genes that are clustered in one species are often reordered in another (2,4). For example, multiple alignment of different strains of *E. coli* and *Yersinia pestis* reveal large changes in gene ordering (synteny) across the chromosome of each species (5,6), where fragments of different length undergo rearrangement due to inversion and translocation events. In addition to variation in ordering, gene content is also variable across strains of the same species. As little as 11% of all *Escherichia coli* genes are present in all strains (7,8) and an even smaller 1% in *Pseudomonas aeruginosa* (9). These ever-present genes are described as the core genome, and the larger set of variably present/absent genes as the accessory genome. Together, the entire set of genes across a species is termed the pan-genome.

The forces that contribute to bacterial genome organization in the face of perturbing mutational pressures have been the focus of extensive debate, leading to a menu of alternate models. The influential selfish operon model (10) views gene clustering as a selfish property of an operon, as clustering of physiologically related units enhances the probability that an entire operon will be horizontally transferred together as a functional unit, and therefore overcome selection for the loss of multi-gene functions that are under weak or fluctuating selection. A central prediction of the selfish operon model is that operons are enriched for peripheral (versus core) metabolic functions subject to fluctuating selection, which is supported by anecdotal reports of accessory functions carried on plasmids (11). Systematic whole-genome analyses, however, have not generally supported this prediction, highlighting a greater frequency of clustering for essential genes in *E. coli* (12,13).

Other models of operon evolution highlight the efficiency gains of sharing a single promotor. For example, by reducing the target size for loss-of-function mutations (14), or by offering more complex temporal co-ordination of co-regulated processes (15,16). Moreover, gene clustering, even without a shared promotor, can offer efficiency gains by spatially co-localizing within the cell gene products that engage in related biochemical tasks (17–19).

In this study we propose that the models above represent special cases of a broader gene-interaction model (e.g., metabolic or regulatory connections between genes). The potential importance of gene interactions has been proposed before in the context of operon evolution (20,21), following Fisher’s (22) foundational work on epistasis, recombination, and the evolution of linkage. Fisher reasoned that physical linkage (clustering) between genes can occur, at higher rates than expected by chance, due to positive epistatic interactions. The relevance of this logic to bacteria has subsequently been questioned (10) due to the generally low rates of homologous recombination in clonally reproducing microbes (2,10,23). Here we return to Fisher’s model and apply it considering our broader understanding of the peculiarities of bacterial genomic organization and processes of change beyond recombination.

Our hypothesis is that gene interactions introduce selective effects on genome-modifying processes which lead to the formation of co-segregating, physically linked clusters of interacting genes. We test this hypothesis via an iterative combination of evolutionary simulation models and pan-genome data. Using closed genome data for 100s of *E. coli* and *P. aeruginosa* genomes, we identify clusters of physically linked and co-segregating genes and show that the resulting co-segregation networks map onto underlying gene-regulatory and metabolic gene interaction networks. The results imply that modular gene interactions are sufficient to guide the evolution of persistent gene clusters and are the primary force shaping genome structural evolution.

## Results

### Simulation models (1): Neutral model of genome evolution

Our overarching hypothesis is that gene interactions ultimately drive the organization of genomes into modular clusters of physically linked genes. In the following sections, we develop a model of pan-genome evolution, based on interactions that impact bacterial fitness. First, we establish a baseline neutral model case where there are no interactions (or where gene interactions have no impact on fitness). In this simplified world, genes still experience chromosomal structure, with genes arranged on a circular chromosome. In this neutral world, it is still intuitively reasonable to expect genome structure to persist as a result of initial conditions: genes that are initially close together will tend to stay close together, as they experience a reduced likelihood of gene separation via recombination or other genome modifying functions.

To test this neutral (no selection) hypothesis of genome inertia, we conduct *in silico* experimental evolution on 100 interacting lineages of bacteria (see methods and supplementary information for details). In brief, each lineage has an average genome size of 2000 genes, and is subject to genomic perturbations of gene gain, loss, and gene rearrangement (via inversion/translocation). Gene gain for a focal lineage is sourced from the pan-genome (the genomes of all lineages), therefore linking lineages together via horizontal gene transfer. Our simulation tracks time in fixation events (e.g., fixation within 1 lineage of a gene-loss event).

In our first simulation we track a single gene pair (*x, y*) that are initially adjacent in 10% of all lineages (absent in other lineages) and follow their resulting organization (physical linkage within genomes and co-segregation across genomes). In figure 1A we track linkage via the average chromosomal distance *d* (the number of intervening genes on the shortest chromosomal path between *x* and *y*, averaged across all genomes where both genes are found). Initially, *d* is zero, reflecting our initial conditions focused on two adjacent genes. Figure 1A illustrates that while linked genes can persist in a state of close linkage for 100s of fixation events, given sufficient time (note log scale on *x* axis), genetic perturbations will eventually separate two focal genes. In Figure 1B we see that by 10,000 fixation events, the distribution of *d* approaches a uniform distribution (Figure 1B), consistent with a simple diffusion model of genetic drift.

**Figure 1.**
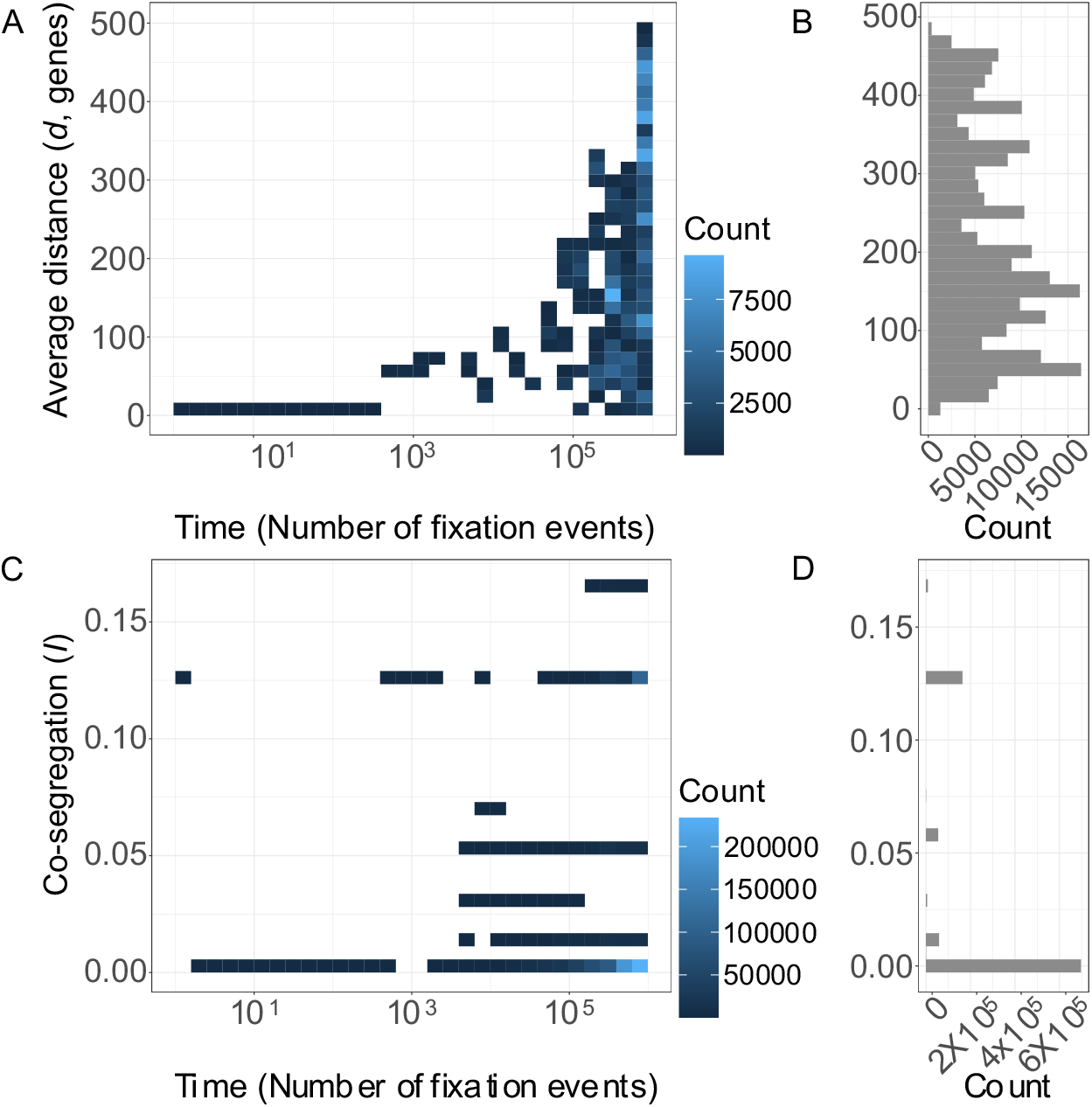
Neutral genomic evolution does not lead to stable co-segregation or clustering in an evolutionary simulation model. An evolutionary simulation of 100 lineages (average 2000 genes per lineage) undergoing neutral events of gene gain and loss and rearrangement. **A.** Average distance *d* between two initially adjacent genes *x* and *y (d*(0) = 0) through simulation time (in fixation events, note log time-scale). **B.** After ∼5×104 fixation events, the distribution of distance *d* approaches a uniform distribution **C.** Co-segregation between a pair of genes in the absence of selection goes down (*I* (0) = 0.127) and fluctuates near a mode of zero. D. After 10^3^ fixation events, modal co-segregation (*I*) = 0.

The distance metric *d* is only applicable for lineages that contain both genes *x* and *y.* Initially, *x* and *y* were always found together (in 10% of lineages; both absent elsewhere) yet processes of gene gain and loss could potentially alter this tight pattern of co-segregation. To assess the extent to which a pair of genes co-segregate, we next estimated the mutual information (*I*) between genes *x*, *y* across the pan-genome (mutual information is analogous to correlation without the need to use real-valued variables (24,25)). In Figure 1C we see that the initial simulation conditions produce an initial mutual information co-segregation value *I* = 0.127. Our results show that over short timeframes (less than 100 fixation events), initially linked genes maintain high mutual information, however as linkage breaks down due to inevitable neutral mutational events, so does co-segregation (Figure 1C; note *I* takes on discrete values due to the discrete nature of presence /absence values used in the pangenome characterization). At equilibrium we observe a skewed distribution of *I* values, with a peak at zero (Figure 1D). Figures 1A-D indicate that under a neutral model, gene pair distance *d* tends to drift and co-segregation *I* peaks at zero.

### Focal gene interaction model

The absence of persistent linkage in our neutral simulation model (Figure 1) is not consistent with examples of highly conserved linkage across bacterial isolates, indicating a role for selection. For example, the *peg* (polyethylene glycol-degradative) operon is conserved across multiple lineages of *Sphingomonas* (26), likewise several *E. coli* operons are conserved in different lineages (27). Therefore, to assess the role of selection in our simulation model, we develop a model to include gene interactions that impact fitness. In this model, events of gene gain, gene loss, and gene rearrangement are now potentially selectively non-neutral if they perturb a gene interaction with defined importance for fitness. As in our previous simulation we track a single pair of genes, but now assume this pair has a positive interaction effect on fitness when found together in a genome (and set all other gene interaction effects to zero). In contrast to Figure 1, we reverse our initial conditions, so that our 2 focal genes are initially not closely linked (Figure 2A) and not co-segregating (Figure 2B).

**Figure 2.**
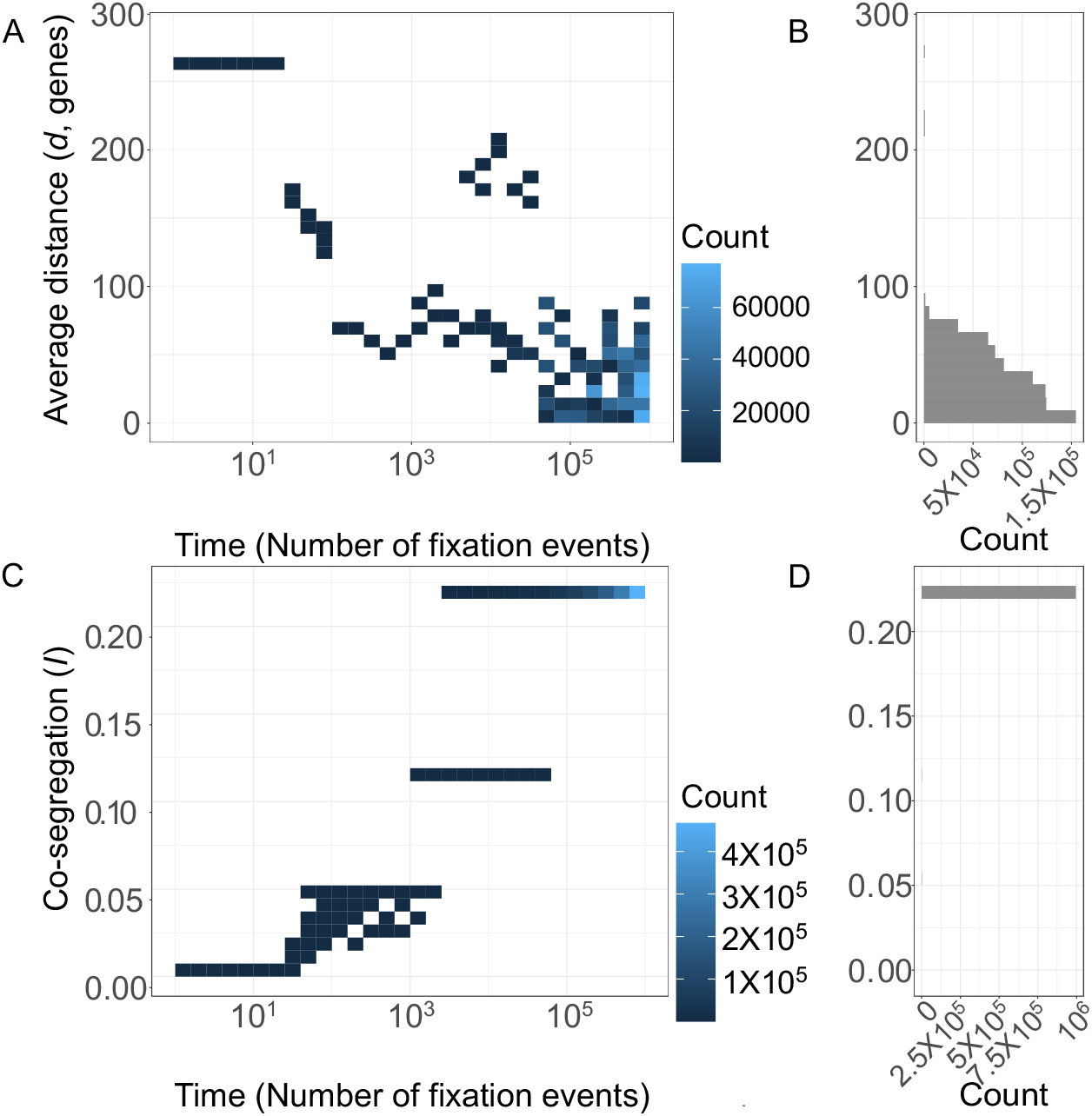
Positive gene interaction leads to close linkage and high co-segregation. An evolutionary simulation of 100 lineages (average 2000 genes per lineage) with apositive defined interaction between two initially non-adjacent genes (*d*(0) = 270). **A.** Average distance between two interacting genes *d* decreases through the course of the simulation and reaches 0 after ∼40000 fixation events (note log scale on x axis). **B.** After 10^6^ fixation events, the modal genetic distance is zero. **C.** Co-segregation *I* between two interacting genes increases during the simulation (from *I*(0) = 0). **D.** After 10^6^ fixation events, modal co-segregation *I* = 0.27.

Figure 2A shows that distance *d* decreases with simulation time, and although stochastic effects can cause instances where distance is high, these are transient events, and the distribution of distances after 200,000 fixation events is stably low, with a mode of zero (Fig 2B). The shortest time to reach a distance of zero (i.e., the focal pair becoming adjacent genes) is ∼40,000 fixation events (Fig 2A) which translates to around 400 fixation events per genome lineage in the collection. Figure 2C shows that during the simulation values of *I* steadily increase. (Because *I* is a global measurement of the linkage of a pair of genes *xy* in the genome collection it is less susceptible to stochastic effects). After ∼15,000 fixation events (150 per genome events) the value of *I* stabilizes around 0.27 and remains the same for the course of the simulation (Figure 2D).

### Genome network interaction model

Figure 2 illustrates that given selection on one gene pair in isolation, close linkage (low *d*) and high co-segregation *I* between these genes can emerge, despite ongoing perturbing forces of gene loss, gain, translocation, etc. Yet in real-world genomes, movement of two positively interacting genes together might in turn pull apart other positively interacting genes. To assess how evolution proceeds given multiple gene interactions, we again extend our simulation model to incorporate *networks* of gene interactions, spanning both random network models (28) and modular interaction network models (small world (29) and preferential attachment (30) models). Specifically, we hypothesize that modular gene interaction networks will drive repeated clustering of interacting genes at a pangenome level.

On a pairwise scale, repeated clustering of interacting genes implies that clustered (low *d*) gene pairs will tend to co-segregate more often (higher *I*), leading to a predicted negative correlation between *d* and *I.* Figure 3A-D plots *d* against *I* under the various interaction models, including the ‘no interaction’ neutral case (Figure 3A). In both the neutral and random interaction case we can reject the hypothesis of a negative association between *d* and *I* (neutral model: *r =* -2.3X10^-3^, *p =* 0.75; random interaction model, *r* =-1.4X10^-3^; p = 0.84). In contrast, when we introduce modular networks, we find support for a negative association (small world model: *r* = -0.05; p = 6.2X10^-3^; preferential attachment model: *r* = -0.12; p = 0.01).

**Figure 3.**
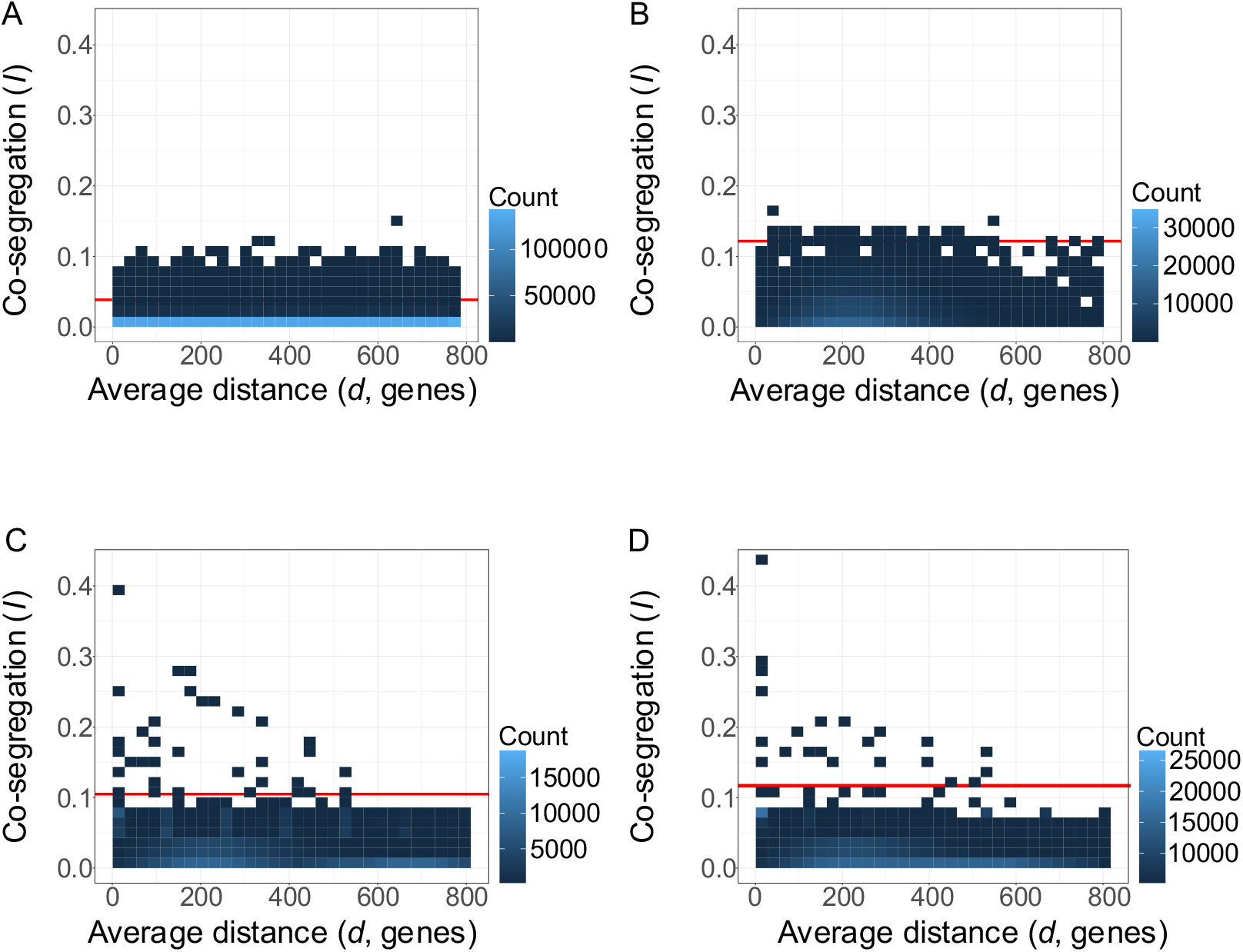
Modular gene interaction networks generate negative associations between chromosomal distance *d* and co-segregation *I.* Values for co-segregation and distance are calculated for all pairs of genes in a collection of 100 genomes with an average genome size of 2000 genes. Threshold of outlier values of *I* is denoted in red (calculated with MAD (31)). **A.** Neutral model. Values of co-segregation for all pairs are lower than 0.15 and distances are uniformly distributed (*r =* -2.3X10^-3^, p *=* 0.75; outlier *I r* = -3.4X10^-4^; p = 0.97). **B.** Interaction effects distributed according to a random network produce no significant linear correlation (*r* = -1.4X10^-3^; p = 0.84; outlier *I r* = -4.1X10^-3^; p = 0.67). **C.** A small world network produces a negative correlation (*r* = -0.05; p = 6.2X10^-3^; outlier *I r* = -0.33; p = 2.6X10^-12^). **D.** A preferential attachment model produces a negative correlation (*r* = -0.12; p = 0.01; outlier *I r* = -0.62; p = 5.2X10^-16^).

While the modular simulations (Figures 3C, D) reveal significant statistical support for a negative association, the effect size is relatively weak and the distribution of simulation results appears to be sparse for higher values of *I*, indicating a potential problem of under-sampling. We note that panels 3C, D summarize the behavior of approximately 10 million gene pairs, so we are not under-sampling. We conjecture that the sparse boundary reflects that the simulated data is a composite of a large ‘background’ of non-interacting pairs producing low *I* values (see neutral simulation case, Figure 3A) overlaid with a smaller set of interacting gene pairs that are biased towards high *I* and low *d* values (Figure 2). Consistent with this conjecture, the gene interaction networks were all sparse (around 0.1% of the possible interactions are included in the network). To further test this conjecture, we identify values of *I* that are outside of the central distribution of *I* values (‘outlier *I* values’) and repeat our statistical tests of association between *d* and outlier values of *I*. We then go on to assess whether outlier values of *I* can be used to recover the underlying architecture of gene interactions in our defined simulation system.

To isolate ‘outlier’ values of *I* from a background of neutral interactions we use mean absolute deviation (MAD) to establish a threshold above which values of *I* are to be considered outliers (red lines, Figure 3). We find that for our neutral model (Figure 3A) and our random network model (Figure 3B) outliers have no correlation with distance (*r* = -3.4X10^-4^; p = 0.97 and *r* = -4.1X10^-3^; p = 0.67 respectively). Whereas in all models of modular gene interaction we find a significant and more substantial negative correlation between outlier values of *I* and *d*. (Small world network, Figure 3C: *r* = -0.33; p = 2.6X10^-12^; Preferential attachment network, Figure 3D: *r* = -0.62; p = 5.2X10^-16^).

### Gene co-segregation network can recover the underlying defined ‘ground truth’ gene interaction network in our simulation model

Our pan-genome evolutionary simulations with modular gene interactions (Figures 3C,D) indicates that evolution will produce networks of co-segregating (high *I*) and physically clustered (low *d*) genes. Following our arguments above, we hypothesize that the networks of co-segregating genes will recapitulate the underlying modular gene interaction networks, which in this simulation world are defined and accessible as a ‘ground truth’ case.

To test this hypothesis, we first recovered the underlying gene interaction network over which the fitness effects of our simulation are based on (our ground truth). Next, we built a network based on the outlier *I* values of co-segregation (the co-segregation network, see methods and supplementary information). In both networks nodes represent genes, and edges represent interaction effects. To measure similarity between the two networks we used Hamming distance (i.e., number of changes a network must undergo to be identical to the other; vertical blue lines in Figure 4). To place these similarity metrics in context, we asked how often we would see the same degree of network similarity (Hamming distance), given a randomly re-sampled gene interaction network. By re-sampling randomly 1000 times from networks with the same underlying algorithm and parameters as the specific ‘ground truth’ case (see methods and supplementary information) we are able to assess the extent to which the observed network similarity diverges from chance expectations. Figure 4 shows that modular gene interaction networks such as small world or preferential attachment produce co-segregation networks that recover with high fidelity the ground truth network (particularly for preferential attachment, where the observed Hamming distance is in the 1^st^ percentile of the reference distribution). In contrast, a random gene-interaction network does not produce a discernable signal in the resulting co-segregation network. The results in Figure 4 indicate that outlier co-segregation values can be a reliable predictor of underlying gene interactions, given a modular organization of gene interactions. We further assess the extent to which repeated simulations identify the same outliers. By fixing the ground truth network and repeating the evolutionary simulation 100 times we find that the repeatability of ‘ground truth’ discovery is maximal, given modular fitness networks (Supplementary Table S1).

**Figure 4.**
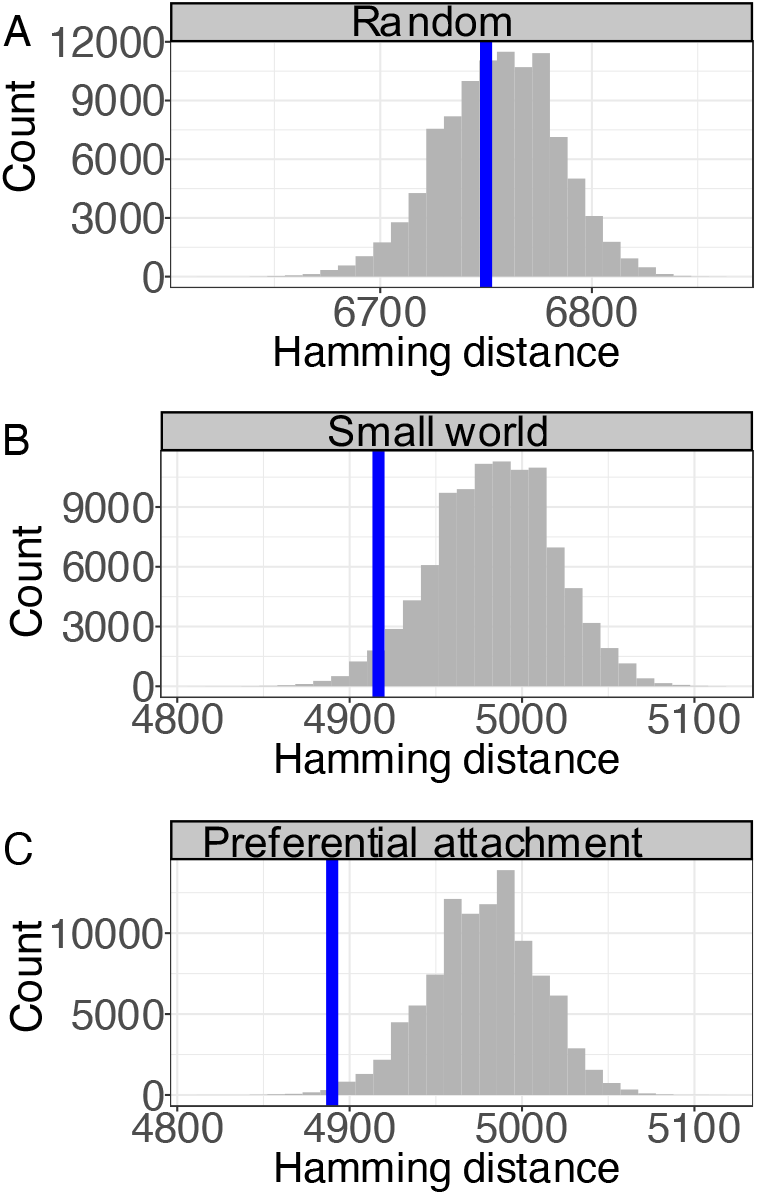
Co-segregation networks identify ‘ground truth’ gene interaction effects in simulated modular networks. Hamming distances for three different gene interaction network algorithms (ground truths) compared to the simulated values of the co-segregation network (network of outlier values of *I*). Blue lines represent Hamming distances between co-segregation networks and ground truth gene interaction networks. Histograms represent distributions of Hamming distances for randomly sampled gene interaction networks (same number of nodes and edges, see methods and supplementary information). A. Random gene interaction networks show distance close to the 50^th^ percentile compared to the background distribution. B. Small world gene interaction network has a distance in the lower 10^th^ percentile of the background distribution. C. Preferential attachment gene interaction networks show a distance within the 1st percentile compared to the background.

### Gene clusters

Our simulation results to date examine pairwise measures of genome structure (pairwise distance *d* and pairwise co-segregation *I*), which does not directly capture the extent of larger-scale gene clustering. In a final simulation analysis, we define a gene cluster as a set of genes that are *repeatedly* adjacent to each other, across a set of genomes. In Figure 5 we plot distributions of cluster sizes, as a function of different threshold values for ‘repeatability’ (defined by prevalence – i.e., the fraction of the genomes in which a cluster is observed). In the absence of gene interactions, we find no clusters larger than gene pairs, even with a permissive prevalence cuttoff of 40% (Figure 5A). In contrast, all our simulations with gene interaction effects show cluster sizes following a geometric distribution, with abundant small clusters and rare large clusters (Figure 5B-D). Unsurprisingly, when gene interactions follow a random network model, clusters are small (Figure 5B). In contrast, small world and preferential attachment networks can lead to bigger cluster sizes (Figure 5C, D).

**Figure 5.**
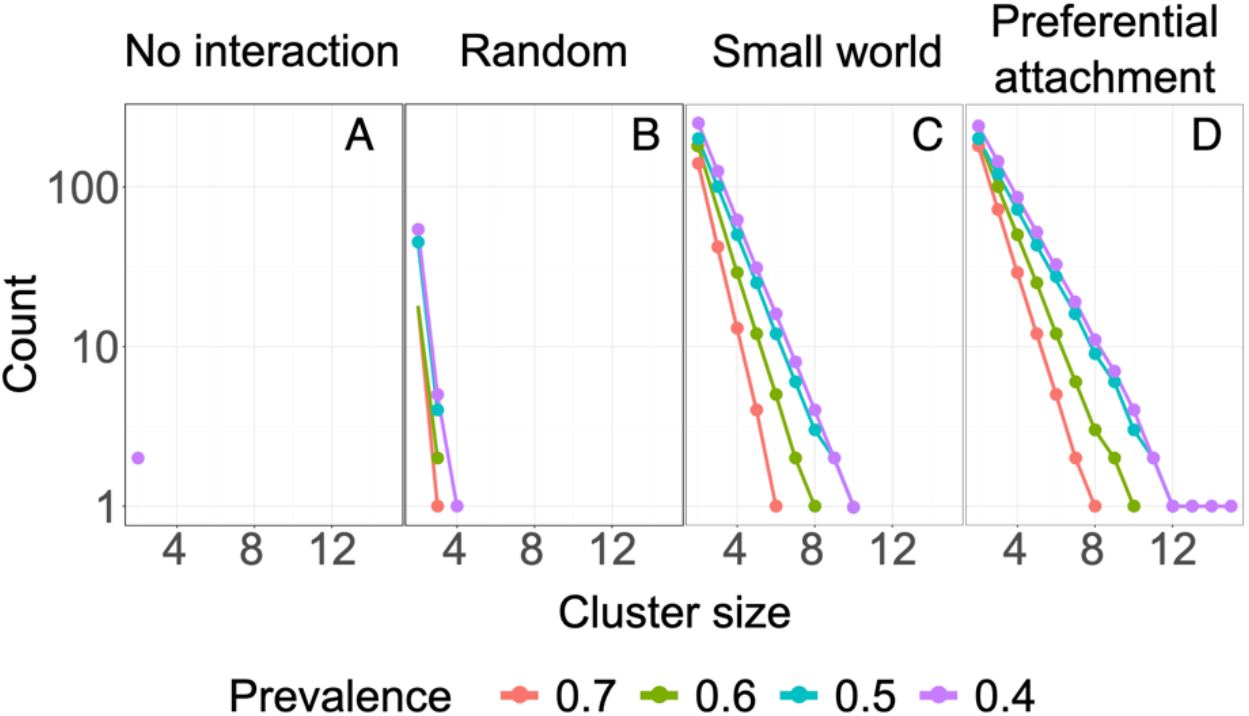
Modular gene interaction networks generate persistent large clusters in bacterial genomes. After evolutionary simulations reached 1 million fixation events, the number of clusters of each size is calculated for four models of gene interaction (**A**, neutral; **B**, random network; **C**, small world network; **D**, preferential attachment network). Clusters are defined as sets of genes that are repeatably observed across the simulated pan genome. Colored lines represent different prevalence thresholds for cluster definition (present in 40% to 70% of all genomes). Preferential attachment and small-world networks yield more clusters and larger cluster sizes.

### Model assessment using pan-genomes of closed P. aeruginosa and E coli genomes

Our simulation models produce several predictions that we now test using genome data on the model species *Escherichia coli* and *Pseudomonas aeruginosa*. We examine genetic linkage, cluster formation and their relationship to genetic networks to evaluate our model predictions.

To capture information on chromosomal linkage, our analyses require the use of completed genomes only. Our results are based on 329 *E. coli* and 179 *P. aeruginosa* closed genomes from the NCBI database that met our criteria for inclusion (see methods and supplementary information). For each individual genome we predicted the coding sequences (CDS), finding on average 5654 CDS for *E. coli* and 6159 for *P. aeruginosa*. We then compared the predicted CDS using BLAST reciprocal best match to determine orthologous groups (OGs, i.e., genes with a common evolutionary origin within a species) with Markov clustering. Given our focus on completed genomes only, the estimated pan-genomes (38731 OGs across all genomes of *E. coli* and 16795 for *P. aeruginosa*) are smaller than estimates using all available sequence data (9,32). Because most of the genes are found in the accessory genome, a matrix of presence/absence of genes in each genome is highly sparse and bears the characteristic histogram U shape (Figure S1, Baumdicker, Hess, & Pfaffelhuber, 2012; Collins & Higgs, 2012; Lobkovsky, Wolf, & Koonin, 2013). A key aspect of our analysis is to examine the extent of co-segregation among gene pairs, which we can only calculate for variably present accessory genes. We therefore focus in this study on the accessory genome of *E coli* (5035 genes) and *P. aeruginosa* (4973 genes).

### Testing cluster formation predictions

Our previous simulation results (Figure 5A) predict that under the neutral model clusters of size 2 are rare and larger clusters are effectively absent. To quantify the proportional abundance of clustered genes in the neutral simulation model, we find that 0.02% of the pan-genome are found in persistent gene clusters of size 2 or more (measured with prevalence cutoff of 0.4, purple line in Figure 5A). In contrast, the selection models allow for larger and geometrically distributed clusters, particularly under modular interactions. Specifically, we found the proportion of clustered genes to be 0.9%, 6% and 8% under the random, small-world and preferential attachment models, respectively (Figure 5B-D).

To assess these predictions against data, we examine the cluster distribution in our pangenome data collections, using the same prevalence thresholds and defining adjacency as a set of genes that are within 10KB of each other. Like our simulated results, cluster size approximates a geometric distribution (Figure 6A). Indicating that large gene clusters although present are rare in nature. Combining large and small clusters together we find that the total fraction of persistently clustered genes (6% for *E. coli* and 12% for *P. aeruginosa*) is in agreement with our modular network simulation models. In Figure S2 we examine the sensitivity of our results to the assumption of 10KB proximity, illustrating that cluster formation follows a geometric distribution in all cases (the number of clusters and maximum cluster size is lower when the proximity window is smaller).

**Figure 6.**
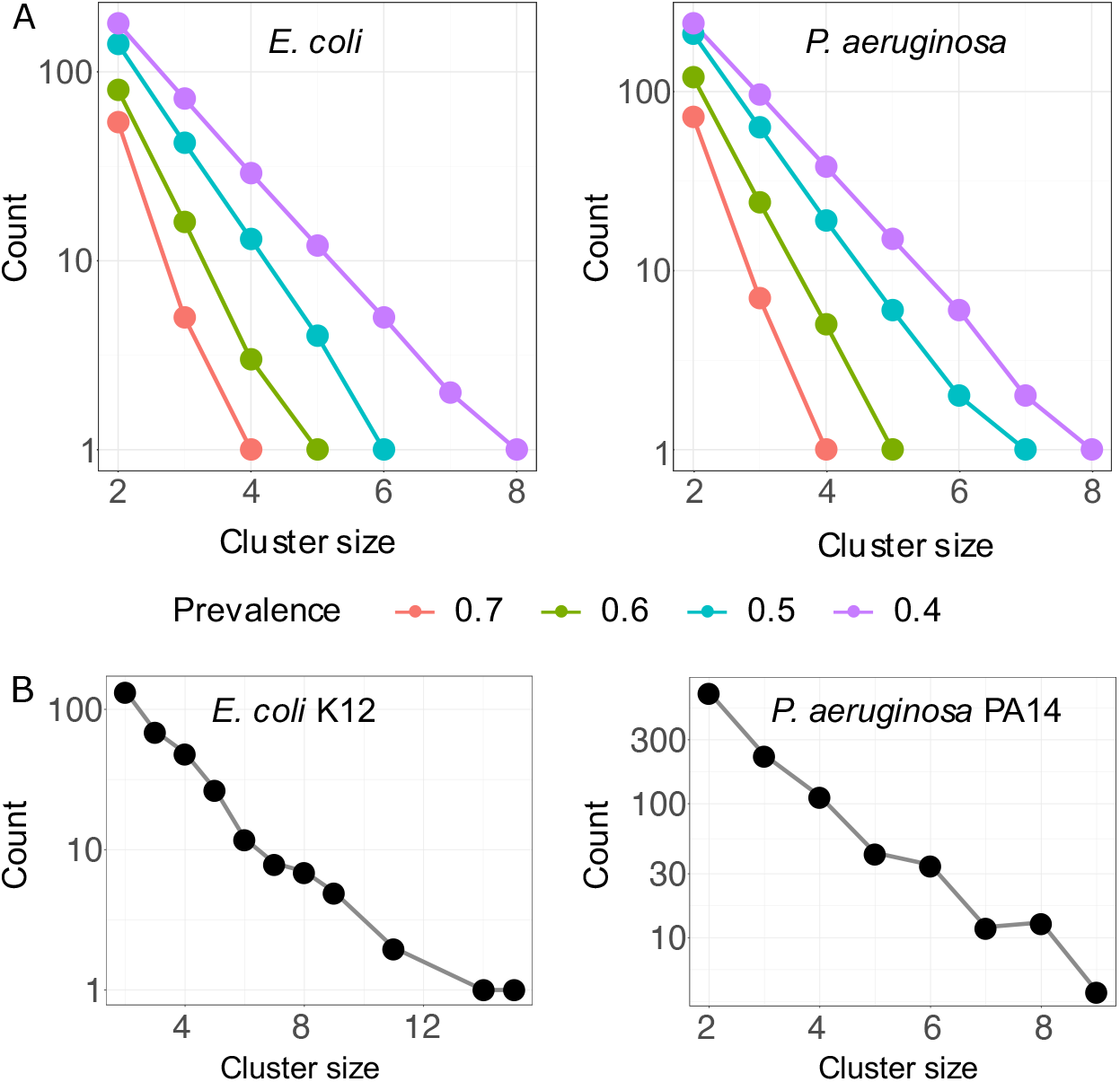
Persistent gene clusters are distributed geometrically across the pan-genomes *E. coli* and *P. aeruginosa*. **A.** Clusters of genes in the *E. coli* and *P. aeruginosa* pangenome, lines represent different prevalence for clusters in the genome collection. In both cases data follows a geometrical distribution like the simulation results. **B.** Defined operon cluster sizes from model strain genomes are distributed according to a geometric distribution for both *E. coli* and *P. aeruginosa*.

Next, we used experimentally validated operons as a reference for the distribution of cluster size in a single genome. We consulted the RegulonDB for *E. coli* (36,37) which includes experimentally validated operons for the strain K12. The study on the transcriptome of *P. aeruginosa* by Wurtzel *et al*, (38) was used to estimate operons of genes in the strain PA14. In the latter an operon is defined as a set of genes transcribed together. In both cases the distribution of sizes approximates a geometric distribution (Figure 6B) in agreement with a modular network model (Figure 5C, D).

### Co-segregation versus linkage

Figures 3 C, D predict a negative relationship between *d* and outlier values of *I*, given modular gene interactions. Turning to our genomic data (Figure 7), we see a predominance of low *I* values, regardless of chromosomal distance *d*. A simple correlation of *I* and *d* values reveals a significant yet small negative association (Figure 7A *E. coli:* Pearson correlation *r =*-0.02; p *=* 0.003. (Figure 7B *P. aeruginosa: r =* -0.012; p = 0.001). To analyze the associations with *d* for outlier values of *I,* we proceed with the same data analyses as in our simulation models. After identifying outliers via the median absolute deviation (31) we find a more substantial negative association between outlier *I* and *d* values (Figure S3; *E. coli: r = -*0.34; p *=* 2.2X10^-6^*. P. aeruginosa: r =* -0.23; p *=* 1.5X10^-6^), consistent with our modular network simulation results. Figure 7 shows a pattern of sparse outlier values of *I* that resembles our modular interaction models (Figure 3C, D), demonstrating a bias towards the highest measures of co-segregation all occurring among genes in close association (see Figure S3 for a higher resolution plot of this region). Our statistical analysis of Figure 7 data treats values of *d* and *I* as statistically independent, however the underlying gene pair data is derived from bacterial genomes that show variable degrees of recent common ancestry. To address the potentially confounding impact of phylogenetic distance, we repeated our analysis controlling for the effect of phylogenetic relationships. Specifically, for each gene pair we calculated the average phylogentic distance among strains that share this gene pair and added the distance metric as a model covariate (see methods and supplementary information for details). Correcting for phylogenetic distance produces only minor changes to our results. *E. coli,* all data: Pearson correlation *r =*-0.02; p *=* 0.005. *P. aeruginosa all data: r =* -0.008; p = 0.0001. *E. coli,* outlier *I* values*: r =* -0.32; p *=* 1.5X10^-6^. *P. aeruginosa* outlier *I* values*: r =* - 0.19; p *=* 5.6X10^-5^).

**Figure 7.**
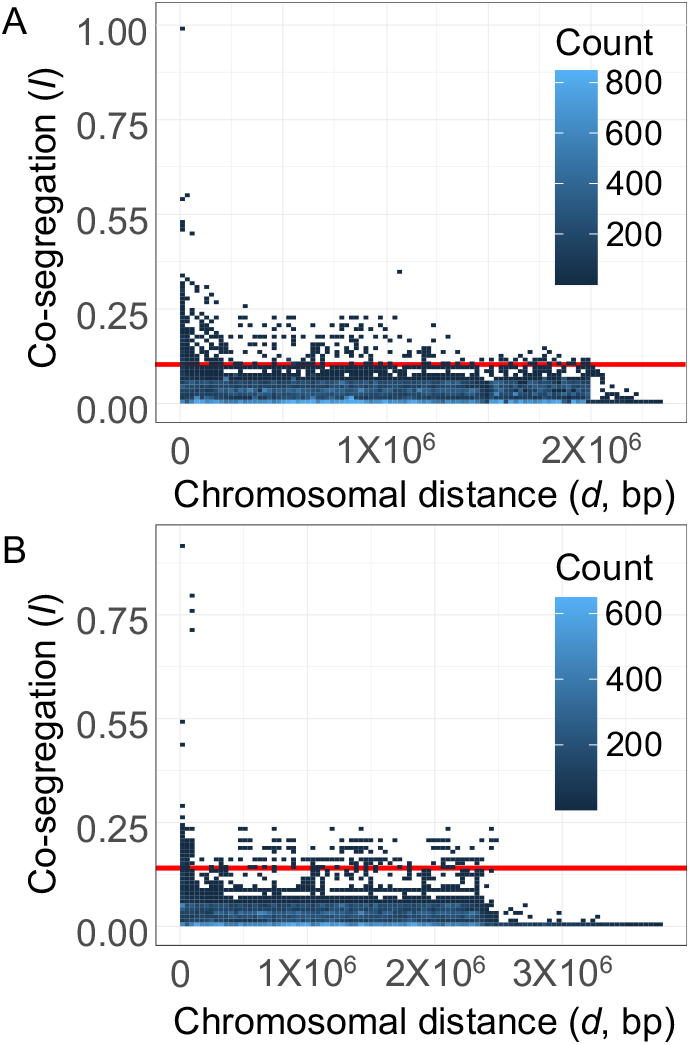
Outlier values of co-segregation *(I)* are negatively correlated with chromosomal distance (*d*) across *E. coli* and *P. aeruginosa* pan-genomes. Each data point refers to an individual gene pair from among the 5035 accessory genes of *E. coli* (A) or the 4973 accessory genes of *P. aeruginosa* (B). For each gene pair *(x,y)*, we calculate average chromosomal distance *d* (across all instances of co-localization within a genome) and co-segregation *I*. Across all gene pairs, we find a small negative correlation between *I* and *d (E. coli: r* =-0.02; p *=* 0.003*. P. aeruginosa: r =* -0.012; p =0.001). Across gene pairs with outlier values of co-segregation (values above the red line: *E. coli: I >* 0.093. *P. aeruginosa*: *I* > 0.012; see Figure S3) we find a stronger negative correlation (*E. coli: r* = - 0.34; p *=* 2.2X10^-6^*. P. aeruginosa: r* = -0.23; p = 1.5X10^-6^).

### Defining the underlying gene interaction mechanism in E. coli and P. aeruginosa

In light of the simulation benchmarking results in Figure 4, we next turn to our measured outlier *I* values for *E. coli* and *P. aeruginosa* and ask: what is the network structure of observed *I* values, and does this structure recapitulate candidate gene-interaction networks, namely published metabolic networks and regulatory networks for these two model organisms?

In our simulation models we had defined ‘ground truth’ gene interaction networks (Figure 4). In our empirical analysis, we turn to candidate interaction networks, beginning with established gene regulatory networks for *E. coli* and *P. aeruginosa* (39–42), involving 1258 accessory genes from *E. coli* and 987 for *P. aeruginosa*. We next assess the similarity of this network with our co-segregation network via Hamming distances, and again contrast this observed network distances with randomized gene interaction network expectations (Figure 8A, B). The Hamming distance to the regulatory network is in the 1^st^ percentile for *E. coli* and 25^th^ percentile for *P. aeruginosa* (Fig 8 A, B).

**Figure 8.**
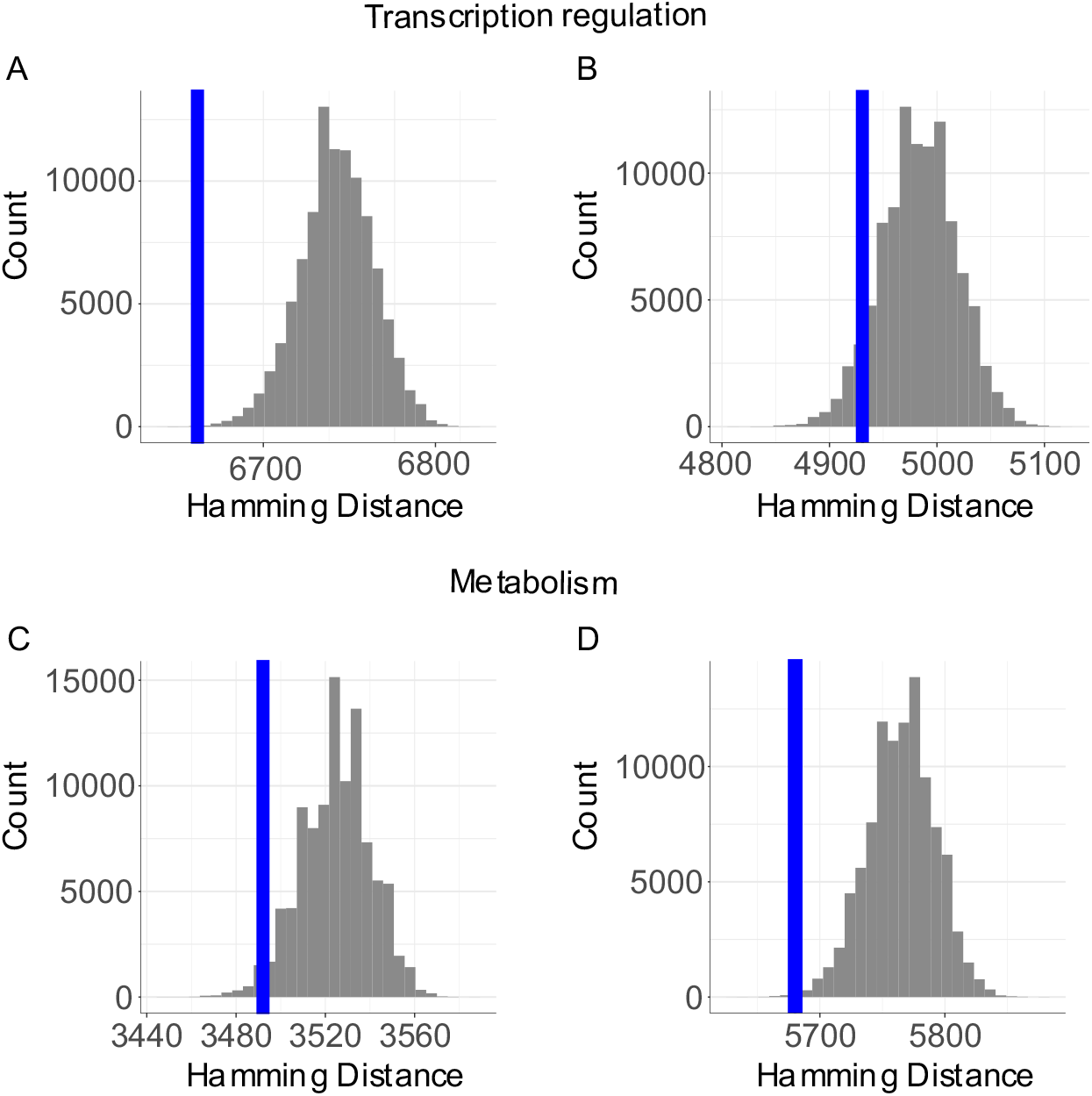
Co-segregation networks capture properties of both regulatory and metabolic networks in bacteria. Hamming distance between a co-segregation network obtained from the pan-genomes of 329 genomes from *E. coli* (panels A, C) and 179 genomes of *P. aeruginosa* (B, D) and their respective transcription regulatory network (A, B) and metabolic networks (C, D). Histogram data represents the distribution of comparisons of 100000 bootstraps of the networks. The vertical blue line represents the measured value. A, B. Assessment of regulatory network for *E. coli* and *P. aeruginosa* respectively. C, D. Assessment of metabolic networks for *E. coli* and *P. aeruginosa* respectively.

In addition to evaluating regulatory networks, we performed the same analyses for established metabolic networks for both species (43,44), which contain 1538 accessory genes of *E. coli* and 1278 genes of *P. aerugiunosa*. In this case we consider an edge between two genes if they participate in the same metabolic reaction or an adjacent reaction (i.e., the products of one reaction are reactants in the adjacent). We observe the Hamming distance for *E. coli* to be around the 10^th^ percentile and less than the 1^st^ percentile for *P. aeruginosa* (Fig 8 C, D).

To further test these conclusions, we assessed the extent to which the entire co-segregation network (not just outlier values of *I*) can serve as a predictor gene interaction network structure, for both metabolic and regulatory gene interaction networks (AUROC analyses, Supplementary Figure S4 A, B). Consistent with Figure 8, we find that the *E. coli* co-segregation network is most predictive of the *E. coli* regulatory gene network (AUROC = 0.86) and the *P. aeruginosa* co-segregation network is most predictive of the *P. aeruginosa* metabolic network (AUROC = 0.96). Altogether, the results in Figures 8 and S3 show that co-segregating genes in the accessory genome are more closely associated with direct regulatory links in *E. coli* (Figure 8A, S3A) and direct metabolic links in *P. aeruginosa* (Figure 8D, S3D). The genomic data on co-segregation networks underlying Figures 7 and 8 is available as Supplementary Table S2 and via the Brown lab repository (https://github.gatech.edu/jcastro37/genome_evolution).

## Discussion

Our pan-genomic results reveal that co-segregating genes (high *I*) tend to be located closer on the genome (low *d*), for two model bacterial systems (Figure 7). While physical linkage provides an intuitive explanation for this result, our neutral evolutionary simulations show that initial linkage provides only a transient support for this association (Figure 1). Introducing selection on gene interactions into our simulation models can produce persistent linkage and co-segregation for a focal gene pair (Figure 2). When gene interactions are distributed modularly across the genome, we see the emergence of persistent clusters of chromosomally linked and co-segregating genes (Figure 3, 5), producing patterns of co-segregation that can reveal the underlying defined gene interaction network (Figure 4). Consistent with our simulation results, we find that *E. coli* and *P. aeruginosa* pan-genomes contain clusters of linked and co-segregating genes (Figures 6,7). Assessing our gene co-segregation networks for these two species against candidate gene interaction models we find support for pan-genome evolution primarily driven by metabolic (*P. aeruginosa*) and regulatory (*E. coli*) gene interactions (Figures 8, S3).

The identification of gene interactions is a major topic in biology and has resulted in numerous algorithms to define interactions from comparative genome data (45–50). A key differentiating factor in our work is the use of bacterial closed genomes, which allows us to analyze the dependency of genome structure on gene interactions. Our results have broad implications in both applied and basic biology contexts. In the context of basic biology, our results indicate how pan-genome structure is causally connected to gene interaction networks. On an applied scale, this causal connection can be used to aid gene function annotation. High co-segregation values can be taken as evidence for positive gene interactions, providing a tool to infer functional links from genes of unknown function to genes that are already annotated (e.g., established regulatory or metabolic functions). This approach can be viewed as an extension to the common ‘guilt by association’ framework (51), now leveraging repeated associations on the pan-genome scale. Current functional network models, including our baseline regulatory and metabolic networks examined above, are heavily biased towards the gene content of model strains(37,42–44,52). Our approach offers a path to infer and extend network structures beyond the well-studied domain of reference genomes.

We note that our results are potentially dependent on several limiting assumptions, for example we currently only examine the impact of positive versus neutral fitness interactions. Although this assumption is sufficient to generate gene clusters in our simulation, avenues for future work could include examination of the effects of negative and positive interactions on cluster formation, and/or higher-order fitness interactions (i.e., pairwise effects dependent on genetic background)(50,53,54). By focusing on orthologous groups of genes, gene-dosing effects of gene duplication are also omitted from our analysis (55,56). Our focus on accessory genomes captures the large majority of the pan-genome of *E. coli* and *P. aeruginosa*, but omits the core genome (1.2 % and 4.5 % of *E coli and P.aeruginosa* pangenomes, respectively). It is reasonable to expect that core genes have interactions with the variable genome which we are not considering.

By modeling time via fixation events, we do not require parameterization of absolute rates of fixation, and therefore of the strength of selection. Our model captures two limit cases for the strength of selection: (1) No selection, where drift is the driving evolutionary force (Figure 1) and (2) selection dominating drift (Figures 2-5). In this second scenario, increasing strength of selection translates to more rapid fixation events, leaving the underlying model behavior unchanged when time is measured in fixation events (Figures 1-5). We do need to make assumptions however on the relative rates of distinct fixation events in our simulations. Following from an assumption of genome size stability (i.e. no directional change in lineage genome size through time, (54,57)), we set rates of lineage gene gain and gene loss to be equal. Following previous work ((58–60)), we further assume that genome rearrangement fixation events are substantially (three orders of magnitude) lower than lineage rates of gene gain and gene loss. While consistent with prior data and simulations, the fixation of genome rearrangement events is reported to vary widely across species (Ballouz et al. 2010). To examine the impact of varying the relative rates of fixation events, we find that our results are robust to order-of-magnitude variation in the relative rate of genome rearrangement fixation events. Specifically, relative rates of between 10^-3^ and 10^-5^ rearrangement fixations per gain (or loss) fixation result in stable cluster formation (Figure S5).

We have shown that gene interactions can introduce systematic biases to pan-genome evolution. In particular, modular gene interactions are sufficient to guide the evolution of persistent gene clusters, and we argue are the first step in the evolutionary process of operon formation. Although several other models have offered working alternatives to the clustering/operon problem, prior work lacks explicit treatment of the pan-genome. By integrating simulation and data analysis at the pan-genome scale, our work indicates that genome interactions are the central organizing principle to understand the origins and maintenance of pan-genome structure and open a new path for gene annotation across the major pool of bacterial genomic diversity – the accessory genome.

## Methods summary

An evolutionary simulation model was created to analyze pan-genome evolution in 100 evolving lineages (linked by horizontal gene transfer) across the course of 10^6^ fixation events. Each genome is an ordered arrangement of OGs, where the distribution of OGs approximates the observed U-shaped distribution in bacterial genomes (61). Simulations span neutral and selection models, where gene interactions are distributed according to one of three network models: Random (28), Small-world (29) or Preferential attachment (30). All three models are parametrized to generate sparsely populated networks (62). Physical distances *d* and co-segregation values *I* (mutual information) are calculated for all pairs of genes during the course of evolutionary simulations. Outlier values of co-segregation are estimated using median absolute deviation (MAD, 38). Model predictions were tested using a collection of complete genomes from *E. coli* and *P. aeruginosa*. OGs were predicted from predicted coding sequences (63), using BLAST reciprocal best matches and Markov Clustering (64, 65). Distances *d* for all pairs of genes were calculated in base pairs (bp) likewise co-segregation *I* was calculated for all pairs. Detailed descriptions of computational models and methods are provided in the supplementary information files.

## Acknowledgments

We thank Peng Qiu, Marvin Whiteley, Gabriel Perron, Tim Read, Rohan Mehta, the Brown lab and members of the Center for Microbial Dynamics and Infection for valuable feedback on the manuscript. We thank the NIH (5R21AI156817-02) and the Cystic Fibrosis Foundation (BROWN21P0) for funding this project.

