## Supplementary text for "Modular gene interactions drive modular pan-genome evolution in bacteria"

**Contents**

1. Supplementary methods
   1.1. Simulation model of genome evolution
        1.1.1. *Simulation outline*
        1.1.2. *Details of model components*
        1.1.3. *Parametrization, sensitivity analyses*
   1.2. *E. coli* and *P. aeruginosa* genome processing
   1.3. Data analysis for simulation and genomic data
        1.3.1. *Co-segregation and chromosomal distance*
        1.3.2. *Outlier detection and network inference*
   1.4. Data and code sharing
2. Supplementary information references
3. Supplementary figures
4. Supplementary tables
5. Supplementary dataset information
1. **Supplementary methods**
   1.1. **Simulation model of genome evolution**
        1.1.1. *Simulation outline*

A collection $G$ of 100 genomes is simulated through evolutionary time (each genome through time forms an evolutionary lineage, coupled to other lineages by processes of gene transfer). For simulation purposes a genome is an ordered set of genes, where each element represents a gene. Each genome has an average size (*S*) of 1000 genes. Each gene is allocated to a set of genomes by sampling a beta distribution, such that the number of times each gene appears in the collection ($N_{g}$) is given by:

$$N_{g}=B\left( \alpha=0.5,\beta=0.5 \right);$$

This distribution is like the one found in *E. coli* and *P. aeruginosa* pangenomes and is common in bacterial pangenomes (1). $G$ has an associated matrix $W$, which describes the interactions between genes. The $W\left( i,j \right)$ element represents the associated cost or benefit to a genome of carrying both gene $i$ and gene $j$. We recover the neutral model (Figure 2) when $W_{ij}$= 1 for all *i* and *j.*

The simulation considers events of gene gain, gene loss, and genome rearrangement (i.e., genomic inversions and translocations) as independent events the genome can suffer, these events can be influenced by a selection criterion (see next section 1.1.2.) or can be free of selection. The model however does not consider the population dynamics that lead to fixation of genomic changes. It is assumed that (given enough time) a genome lineage will suffer one of the following three categories of possible fixation events: gene gain, gene loss or gene inversion / translocation. Each iteration step of the model represents a fixation event for one of these processes in one of the 100 genome lineages. The simulation algorithm can be summarized as follows:

1. Initialize the set ($G,W$) as described above.
2. Chose an individual genome at random.
3. A fixation event occurs:
   i. Gene gain (see section 1.1.2.).
   ii. Gene loss (see section 1.1.2.).
   iii. Gene inversion/translocation (see section 1.1.2.).
4. Distance and mutual information are calculated for the genes in the set (see section 1.3.1.).
5. Repeat steps 2-4 until the end of the simulation.
6. Record the values and exit.

*1.1.2. Details of model components.*

The probability of fixation events (step 3 above) is governed by the matrix *W*. It represents the relative differences in cost/benefit that gene interactions provide to a given genome. For a neutral case all elements of *W* have the same value (*W_ij_* = 1 for all values of i and *j*) therefore no biases occur when selecting for insertion sites or removing genes. For a selection case the values of *W* are set to 1 for (sparse) interacting genes according to a network of interactions (detailed in section 1.1.3) and the remaining values to 0.

For a gene gain event, the available pangenome pool is sampled, the insert size (i.e., number of genes to be inserted) is calculated according to a zero truncated Poisson distribution with λ=1. $W$ is used to calculate the insertion site in the receiver genome. For each joint in the genome (i.e., edge between two genes) a weight is calculated such that

$$A_{ij}=1-\frac{W_{ij}}{S}$$

Where $A_{ij}$ is the weight for the joint between genes $i$ and $j$, $W_{ij}$ is the interaction coefficient between the two genes, and $S$ is the genome size. An insertion site is selected by sampling from the distribution of values $A$. We recover the neutral model (Figure 2) given $W_{ij}$= 1*.*

For a gene loss event, fragment size is selected the same way as with gene gain. A fragment to be removed is selected by calculating its weight, by considering the sum of the weights for the genes in the fragment.

$$R_{ij}=\sum_{i=1}^{I} \sum_{j=1}^{J} A_{ij}$$

The fragment to be removed is sampled form the set of all possible weights *R.*

Finally, for a genome rearrangement event. A fragment to be moved is selected same as for gene loss, and the fragment is inserted into a site selected according to gene gain, fragments size is likewise determined by a zero truncated Poisson distribution with λ=1.

*1.1.3. Parametrization, sensitivity analyses*

For all simulations we used a genome collection of 100 separate genome lineages. 1000 genes (assumed to be orthologous groups) are allocated to the collection such that the probability of each gene (P(*g*)) follows a beta distribution with positive equal shape parameters ($\alpha,\beta$), such that ($P\left( g \right)= Beta\left( \alpha=0.5,\beta=0.5 \right)$. Genome size is initially 2000 genes for all genes. Fixation events rates are defined as: gene gain (*P_g_*), gene loss (*P_l_*), and gene rearrangement (*P_r_*).

$$P_{r}=0.001;P_{g}=P_{l}=\frac{1-P_{r}}{2};$$

By defining $P_{g}=P_{l}$, we ensure that average genome size stays constant on longer time scales (but can also fluctuate around this average).

For our model of interactions, a network is defined for all genes (*G*). We use three network generated models:

-A random network according to the Erdős–Rényi random graph model (2), with edge sampling parameter *p* =0.0125.

-A small-world network according to the Watts–Strogatz model (3), with an expectation of 1.2X10^6^ edges and an average degree (*K*) of 3.

- A preferential attachment network according Barabási–Albert model (4), with a degree distribution parameter ($\gamma$) equal to 3 and a growth rate parameter of 1.2.

This parameter selection ensured we have networks with similar number of nodes and edges such that the simulations are comparable. Because both the Watts–Strogatz and the Barabási–Albert generate sparse networks we use a small value of *p* in our random network. Specifically small world networks require that the number of edges *I* is smaller than the number of nodes (*n*) such that, $n\gg e\gg ln\left( e \right)$ (3). While preferential attachment networks have a distribution of degree following a power law, they generate sparse network with, $n\gg e$ (5).

To calculate cluster size, we selected 4 different prevalence thresholds (0.4, 0.5, 0.6, 0.7) and selected all genes with prevalence equal or higher in our genome collection. For all gene pairs we consider a cluster if the distance between neighboring pairs is at or below our selection criteria (10Kbp for bacterial genomes, 0 genes for simulated genomes). This process is repeated iteratively to identify larger linked clusters (up to a limit of cluster size = 10).

***1.2. E. coli* and *P. aeruginosa* genome processing**

Data collection and quality control. All genomes were pulled from NCBI on Jan 2023, resulting in 1214 closed genomes for *E. coli* and 824 for *P. aeruginosa*. To avoid sampling issues in the gene content analysis only completed genomes were included, this includes sequences labeled as either “Complete genome” (which includes the chromosome and plasmid information) and “Chromosome” in the database. Plasmid genes were removed so that the chromosomal distance metric *d* is defined for all gene pairs. Additionally, we removed any genome that did not meet our quality control thresholds (completeness of at least 95% and maximum contamination of 2%, (6,7)). We found 179 *Pseudomonas aeruginosa* and 329 *Escherichia coli* genomes from the NCBI database that met our criteria for inclusion.

Orthologous group identification. Coding sequences for each genome were predicted using GeneMarkS v.1.11 (8). For each pair of genomes reciprocal protein BLAST v.2.10.1 (9) was performed. Reciprocal best matches are used to determine orthologous groups (OGs) with Markov clustering, using and inflation value of 1.5 (10). We exclude core genes (i.e., OGs present in all the genomes in the collection), and OGs with a single instance in the collection to produce a dataset of accessory genes (the variable genome).

Phylogenetic analysis. Individual amino acid gene alignments for 102 genes essential to bacteria and archea (11) were analyzed using RAxML v8.0.19 (12). For all RAxML analyses, we used a gamma-proteobacteria evolutionary model which uses an aggregate parameter distribution for site-to-site rate variation and the Jones-Taylor-Thornton amino acid substitution matrix with 100 rounds of bootstrapping (13,14). Individual alignments were concatenated using an in-house concatenation alignment script, concatenated genes were used to generate a phylogeny for the evaluated strains. For each gene pair (when possible) the subtree of strains containing both genes was extracted and the average phylogenetic distance (*p*) was used as a covariate for partial correlation test between chromosomal distance (*d*) and co-segregation (*I*) (7).

**1.3. Data analysis for simulation and genomic data**

*1.3.1. Co-segregation and chromosomal distance*

A presence/absence matrix was built for the selected genes (OGs). In this numeric matrix (1 representing presence and 0 representing absence), each gene is treated as a random variable and its probability distribution is given by its prevalence in the collection. For each gene pair mutual information is calculated according to:

$$I_{xy}=\sum_{y\in X} \sum_{x\in X} p\left( x,y \right){log}_{2}\left( \frac{p\left( x,y \right)}{p\left( x \right)p\left( y \right)} \right)$$

Where $X$ and $Y$ are two separate genes, $p\left( x \right)$, $p\left( y \right)$ are the marginal probabilities for gene $X$ and $Y$ respectively (e.g., $p\left( x \right)$probability of finding gene $X$ in a single genome), and $p\left( x,y \right)$ is the joint probability distribution of gene $X$ and $Y$ respectively.

Chromosomal distance (*d*) is calculated between all pairs of selected genes. Such that:

$$d_{xy}=\frac{\sum_{g=1}^{G} minD_{g}\left( x,y \right)}{G}$$

Where $d_{g}\left( x,y \right)$ is the set of 2 distances in the circular chromosome between genes $x$ and $y$ for the $g$th genome and $G$ is the total number of genomes in the collection. For simulated genomes $d_{g}$ is measured in number of genes, for genomic data $d_{g}$ is measured in base pairs (bp).

*1.3.2. Outlier detection and network inference*

We used median absolute deviation (MAD) to determine outlier measurements of co-segregation according to (15):

$$MAD\left( I \right)=\mathrm{median}\left( \left| I_{i}-\mathrm{median}\left( I \right) \right| \right)$$

Where *I* is the set of measurements of co-segregation. A given value is considered an outlier if it has an absolute deviation from the maximum deviation larger than 2.

A network is inferred using outlying values of co-segregation. Where for a gene pair *x* and *y,* an edge is drawn between them if their co-segregation (*I*) is larger than the MAD cutoff. Comparison between networks is performed by calculating the Hamming distance that is; the number of transformations (i.e., edge additions and subtractions necessary for the two networks to be identical, this is expressed as:

$$H_{ij}=\left( E\left( G_{i} \right)\cup E\left( G_{j} \right) \right)-2E\left( G_{i}\cap G_{j} \right)$$

Where $E\left( G_{i} \right)$ is the set of edges in network $i$, $E\left( G_{j} \right)$ is the set of edges in network $j$.

**1.4. Data and code sharing**

Source code for the pangenome evolution simulation is available under GNU General Public License v3.0 at GitHub (http://gatech.github.com/jcastro37/genome_evolution). The simulation requires Julia version 1.0.3, or later. Supplementary dataset S1 includes tables with values for distance *d*, co-segregation *I* and average phylogenetic distance for the genome collections of *E. coli* and *P. aeruginosa*. Supplementary dataset S2 includes fasta files with representative nucleotide sequences for the orthlogous groups of both *E. coli* and *P. aeruginosa* genome collections. Supplementary datasets are also available at GitHub. Supplementary dataset S3 includes with data for presence/absence foe each OG in the genomes used for the genome collection of *E. coli* and *P. aeruginosa* including.

2. **Supplementary information references**

1.	Lapierre P, Gogarten JP. Estimating the size of the bacterial pan-genome. Trends Genet. 2009 Mar;25(3):107–10.

2.	Erdos P, Rényi A. On the evolution of random graphs. The Structure and Dynamics of Networks. 2011 Oct 23;9781400841356:38–82.

3.	Watts DJ, Strogatz SH. Collective dynamics of ‘small-world’ networks. Nature. 1998 Jun 4;393(6684):440–2.

4.	Albert R, Barabási AL. Statistical mechanics of complex networks. Rev Mod Phys. 2002 Jan;74(1):47–97.

5.	Genio CI Del, Gross T, Bassler KE. All scale-free networks are sparse. 2011;

6.	Parks DH, Imelfort M, Skennerton CT, Hugenholtz P, Tyson GW. CheckM: assessing the quality of microbial genomes recovered from isolates, single cells, and metagenomes. Genome Res. 2015 Jul 1;25(7):1043–55.

7.	Rodriguez-R LM, Gunturu S, Harvey WT, Rosselló-Mora R, Tiedje JM, Cole JR, et al. The Microbial Genomes Atlas (MiGA) webserver: taxonomic and gene diversity analysis of Archaea and Bacteria at the whole genome level. Nucleic Acids Res. 2018 Jul 2;46(W1):W282–8.

8.	Besemer J, Lomsadze A, Borodovsky M. GeneMarkS: a self-training method for prediction of gene starts in microbial genomes. Implications for finding sequence motifs in regulatory regions. Nucleic Acids Res. 2001 Jun 15;29(12):2607–18.

9.	Altschul SF, Gish W, Miller W, Myers EW, Lipman DJ. Basic local alignment search tool. J Mol Biol. 1990;215(3):403–10.

10.	Gibbons TR, Mount SM, Cooper ED, Delwiche CF. Evaluation of BLAST-based edge-weighting metrics used for homology inference with the Markov Clustering algorithm. 2015.

**3. Supplementary figures**

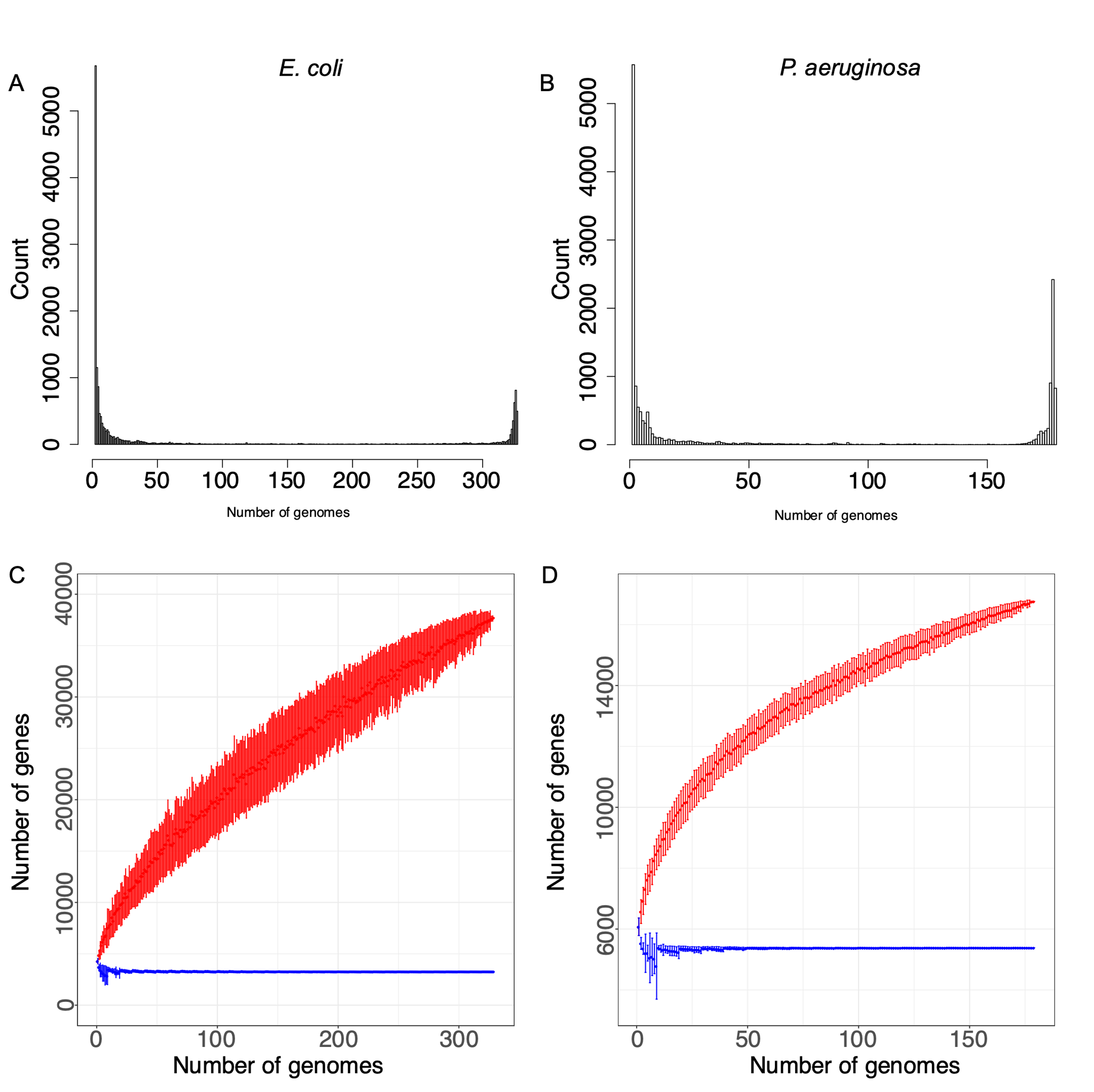

**Supplementary Figure S1. Pangenome sampling of completed genomes of *E. coli* and *P. aeruginosa*.** Identified genes are assigned into orthologous groups via Markov clustering. Distribution of genes across genome collection of 329 *E. coli* **(A)** and 179 P. aeruginosa genomes **(B)**. A significant fraction of genes is present in only one genome, with the majority of genes being present in the variable genome (98% *E. coli* and 96% *P. aeruginosa*). Sampling all genes in the collection is illustrated with red lines, the blue lines denote the core genome. The increasing trend indicates that the pangenome is under-sampled. A total of 38731 genes are identified across all genomes of *E. coli* **(C)** and 16795 for *P. aeruginosa* **(D)**.

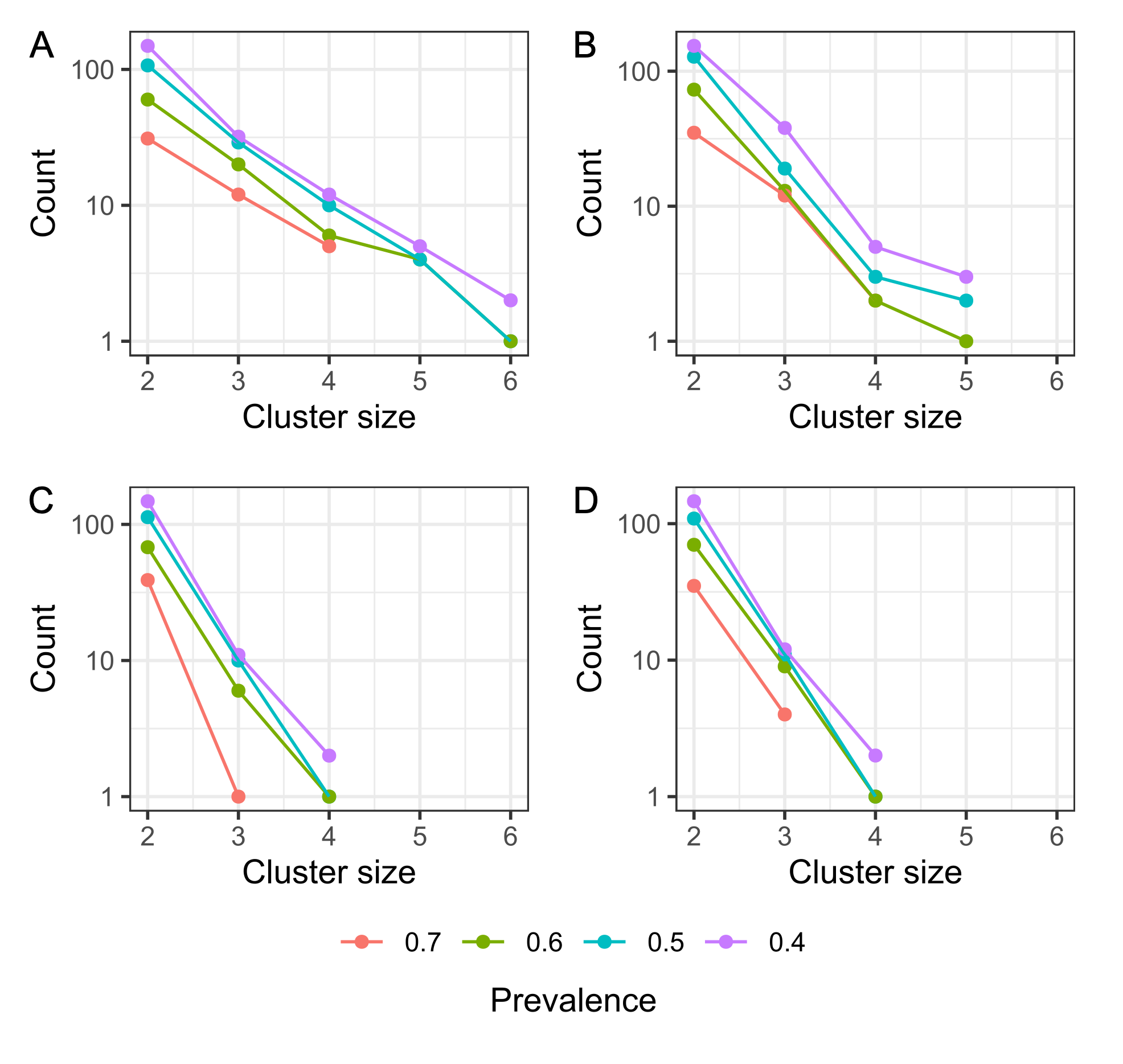

**Supplementary Figure S2. Persistent gene clusters are distributed geometrically across the pan-genomes *E. coli* and *P. aeruginosa*, across different window sizes.** Lines represent different prevalence for clusters in the genome collection. **A, C.** Clusters of genes in the E. coli genome using within a 5kbp window (**A**) and a 2kbp window (**C**). **B, D.** Clusters in the *P. aeruginosa pangenome*, using within a 5kbp window (**B**) and a 2kbp window (**D**). In all both cases data follows a geometrical distribution like the simulation results.

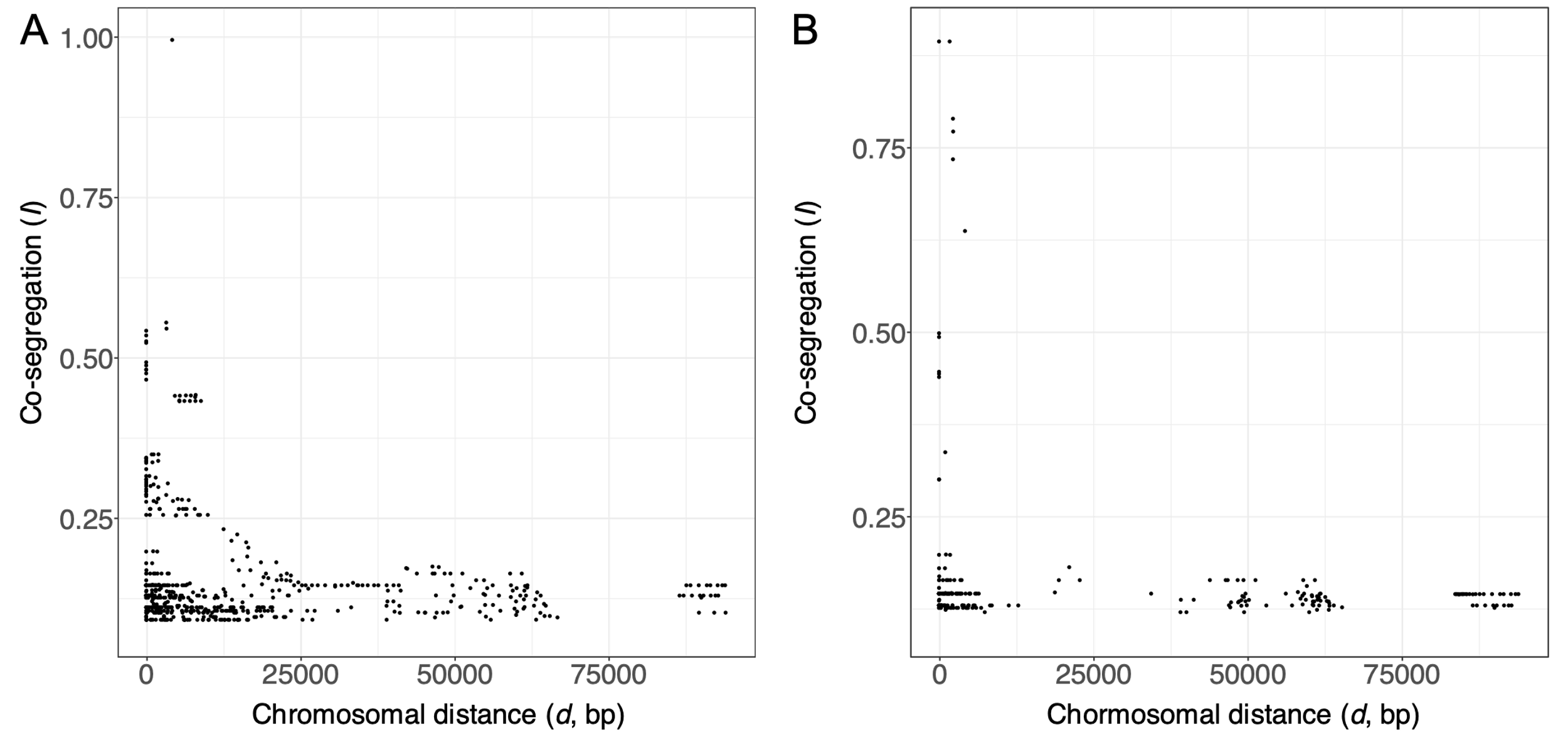

**Supplementary figure S3. Outlier values of co-segregation (*I*) are negatively correlated with distance.** Identified outliers of *I* using median absolute deviation are plotted against chromosomal distance for *E. coli* **(A)** and *P. aeruginosa* **(B)**. In both cases a negative relationship between the two is significant (*E. coli: r* = -0.34*;* p *=* 2.2X10^-6^*.* *P. aeruginosa: r* = -0.23; p = 1.5X10^-6^).

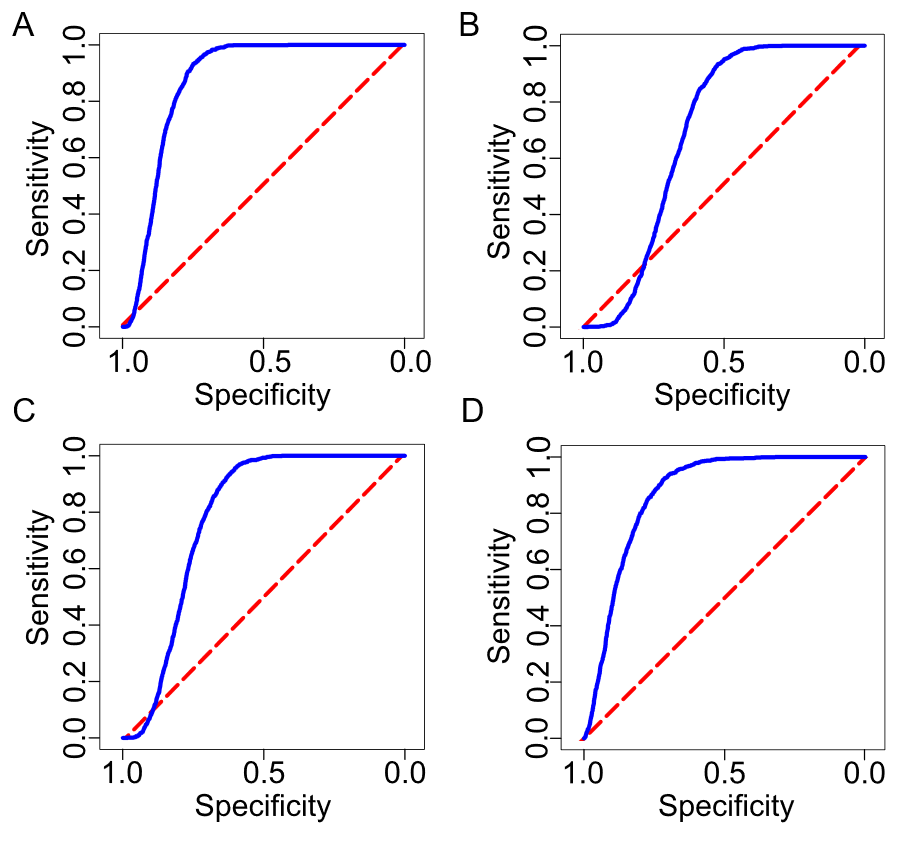

**Supplementary Figure S4. Co-segregation is a competent predictor of regulatory and metabolic interactions in bacteria.** Receiving operating characteristic (ROC) is used as a diagnostic of predictive ability for co-segregation networks obtained from the pan-genomes of 329 genomes from *E. coli* (panels **A, C**) and 179 genomes of *P. aeruginosa* (**B, D**) predicting their respective transcription regulatory network (**A, B**) or metabolic networks (**C, D**). Blue lines represent the predictive ability of co-segregation across a moving threshold in *I*. Red lines represent the expectation of a random guess. **A, B.** Assessment of regulatory network for *E. coli* (Area under ROC = 0.86) and *P. aeruginosa* (Area under ROC = 0.68) respectively. **C, D.** Assessment of metabolic networks for *E. coli* (Area under ROC = 0.78) and *P. aeruginosa* (Area under ROC = 0.86) respectively.

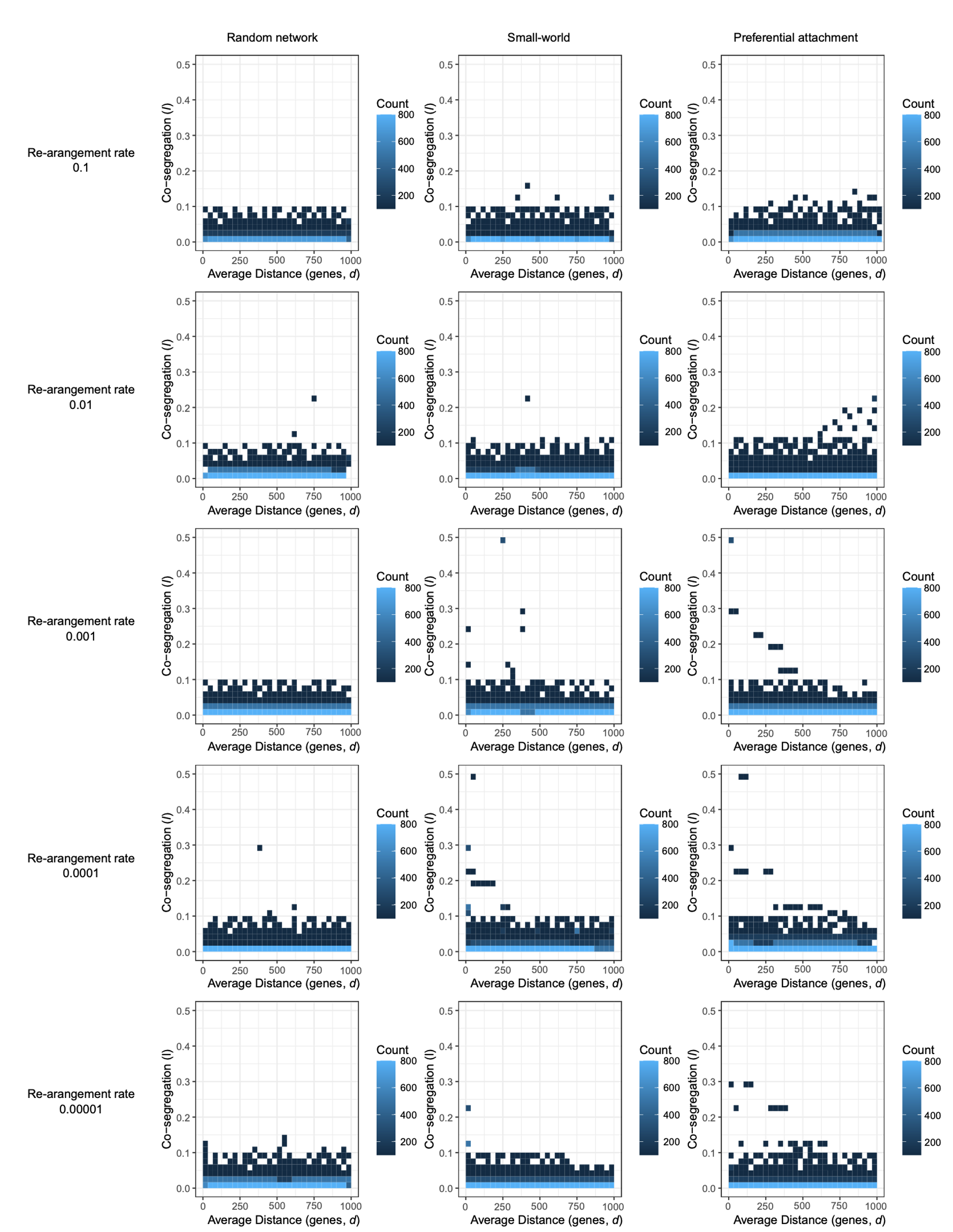

**Supplementary figure S5. The relationship between *d* and *I* is affected by network organization as well as re-arrangement rate due to translocation/inversion events.** A negative relationship can be observed for both small-world and preferential attachment networks, however this relationship is conditional on low re-arrangement rates.

|  | Average network size | Average True Positives | Average False positives | Average AUROC |
| --- | --- | --- | --- | --- |
| Random network | 14994 | 7521 | 7472 | 0.52 |
| Small-world | 15239 | 11416 | 3825 | 0.73 |
| Preferential attachment | 14974 | 11973 | 3001 | 0.79 |

**4. Supplementary tables**

**Supplementary table S1. Outlier values of co-segregation (*I*) can recover the underlying interaction network.** Simulations are repeated 100 times each, in each case the ground truth network and simulation parameters are fixed. Identified outliers of *I* using median absolute deviation (MAD) are used to estimate a network of interaction for the genes in the simulation. This estimate can recover around 50% of the edges in a random network model (the fraction of true positive edges of the total network), 74.9% for a small-world model and 80.0% for a preferential attachment model. Additionally the area under the receiving operating characteristic (AUROC), shows similar values of recovery for all three cases.

**5. Supplementary dataset information**

**Supplementary dataset S1. Values of co-segregation (*I*), chromosomal distance (*d*) and phylogenetic distance (*p*) for gene pairs across *E. coli* and *P. aeruginosa* pan-genomes.** Two datasets are provided, one for *E. coli* (Ecoli_pairwise metrics.tsv) and one for *P. aeruginosa* (Paeruginosa_pairwise metrics.tsv). Each dataset is organized into four columns and xxx / yyy rows (*E. coli* / *P. aeruginosa* respectively). Each row represents metrics for a defined pair of orthologous groups (OG). Column 1 defines the first OG in a pair (OG1), and colum 2 represents the second OG (OG2). colums *d*, *I* and *p* represent values for chromosomal distance (*d*), co-segregation (*I*), and average phylogenetic distance (*p*), for each OG defined pair (each row). A lookup table for locus tags correpsonding to each OG is provided in supplementary dataset S3.

**Supplementary dataset S2. Fasta files with representative nucleotide and amino acid sequences for the OGs in the pangenome of *E. coli* and *P. aeruginosa*.** Each sequence is a representative of the OG, the identifier is the OG id, followed by the genome from which it was pulled and the name of the gene in the individual genome annotation. Ecoli_ogs.fna and Ecoli_ogs.faa contain the sequences for the *E. coli* pangenome for nucleotides and amino acids respectively. Paeruginosa_ogs.fna and Paeruginosa _ogs.faa contain the sequences for the *P. aeruginosa* pangenome for nucleotides and amino acids respectively.

**Supplementary dataset S3. Orthologous group lookup tables for the pangenome of *E. coli* and *P. aeruginosa*.** Tables Ecoli_ogs.tsv and Paeruginosa_ogs.tsv contain data summarizing the presence / absence of each OG in the genomes used in this study. Each row represents an OG and each column a defined genome. The values of each cell are either 0 (absent) or 1 (present). The table Paeruginosa_IS2OGS.tsv includes a relational table of the OG ids used in this study to commonly used locus tags for *Pseudomonas aeruginosa* genomes (as used in the genome database Pseudomonas.com).
